# Recovirus NS1-2 has viroporin activity that induces aberrant cellular calcium signaling to facilitate virus replication

**DOI:** 10.1101/703959

**Authors:** Alicia C. Strtak, Jacob L. Perry, Mark N. Sharp, Alexandra L. Chang-Graham, Tibor Farkas, Joseph M. Hyser

## Abstract

Enteric viruses in the *Caliciviridae* family cause acute gastroenteritis in humans and animals, but the cellular processes needed for virus replication and disease remain unknown. A common strategy among enteric viruses, including rotaviruses and enteroviruses, is to encode a viral ion channel (i.e., viroporin) that is targeted to the endoplasmic reticulum (ER) and disrupts host calcium (Ca^2+^) homeostasis. Previous reports have demonstrated genetic and functional similarities between the nonstructural proteins of caliciviruses and enteroviruses, including the calicivirus NS1-2 protein and the 2B viroporin of enteroviruses. However, it is unknown whether caliciviruses alter Ca^2+^ homeostasis for virus replication or whether the NS1-2 protein has viroporin activity like its enterovirus counterpart. To address these questions, we used Tulane virus (TV), a rhesus enteric calicivirus, to examine Ca^2+^ signaling during infection and determine whether NS1-2 has viroporin activity that disrupts Ca^2+^ homeostasis. We found that TV disrupts increases Ca^2+^ signaling during infection and increased cytoplasmic Ca^2+^ levels is important for efficient replication. Further, TV NS1-2 localizes to the endoplasmic reticulum (ER), the predominant intracellular Ca^2+^ store and the NS2 region has characteristics of a viroporin domain (VPD). NS1-2 had viroporin activity in a classic bacterial functional assay and caused aberrant Ca^2+^ signaling when expressed in mammalian cells, but truncation of the VPD abrogated these functions. Together, our data provide new mechanistic insights into the function of the NS2 region of NS1-2 and show that like many other enteric viruses, enteric caliciviruses also exploit host Ca^2+^ signaling to facilitate their replication.

**Importance:** Tulane virus is one of many enteric caliciviruses that cause acute gastroenteritis and diarrheal disease. Globally, enteric caliciviruses affect both humans and animals and result in >65 billion dollars per year in treatment and healthcare-associated costs, thus imposing an enormous economic burden. Recent progress has resulted in several cultivation systems (B cell, enteroid and zebrafish larvae) to study human noroviruses, but mechanistic insights into the viral factors and host pathways important for enteric calicivirus replication and infection are largely still lacking. Here we used Tulane virus, a calicivirus that is biologically similar to human noroviruses and can be cultivated in conventional cell culture, to identify and functionally validate NS1-2 as an enteric calicivirus viroporin. Viroporin-mediated calcium signaling may be a broadly utilized pathway for enteric virus replication, and its existence within caliciviruses provides a novel approach to developing antivirals and comprehensive therapeutics for enteric calicivirus diarrheal disease outbreaks.

## Introduction

The *Caliciviridae* family consists of small, non-enveloped single-stranded RNA viruses with five major genera: *Sapovirus, Lagovirus, Vesivirus, Nebovirus,* and *Norovirus* (1, 2). Caliciviruses infect a wide-array of hosts and have importance in medical, veterinary, and agricultural fields (3). Of particular importance are human noroviruses (HuNoVs) that are the leading cause of acute gastroenteritis (AGE) in every age group, and can cause life-threatening illness in the young, immunocompromised, and elderly subpopulations (4–7). Estimates show that every individual will experience at least five symptomatic norovirus infections in their life (8), which underlines the need for anti-viral drugs, vaccines, or anti-diarrheal therapies for HuNoV infection (9, 10). However, many aspects of calicivirus pathogenesis, including that of HuNoV, remain uncharacterized, which represents a challenge to developing effective therapies (4, 6, 9). One strategy to address this challenge is to study other enteric caliciviruses, such as porcine sapoviruses and rhesus enteric caliciviruses (*Recovirus*). Recoviruses are a new proposed genus of enteric CVs initially identified in the stool from rhesus macaques, of which Tulane virus (TV) is the prototype strain (11, 12). While Recoviruses constitute a separate genus, these viruses are most closely related to HuNoVs and studies of TV show that it retains both biologic and genetic similarities to HuNoVs, including genomic organization, tissue tropism (intestinal epithelia), and clinical presentation (self-limiting vomiting and diarrhea) (1, 11–13). Furthermore, TV robustly replicates in cell culture in monkey kidney cell lines (e.g., LLC-MK2 cells), which facilitates investigation into the host pathways exploited by TV during infection. This makes TV an excellent model system to identify host signaling pathways broadly exploited by caliciviruses for replication and pathogenesis.

Like other caliciviruses, TV has 3 main open reading frames (ORFs), with ORF1 encoding the nonstructural proteins (NS1-7), and ORFs2-3 encoding the capsid proteins VPI (ORF2) and VP2 (ORF3) (10, 14–15). During replication ORF1 is synthesized into the polyprotein, which is subsequently cleaved by the viral protease NS6 to produce six nonstructural proteins that orchestrate viral replication (4, 12, 14–16). In particular, the roles of the NS1-2 protein (N-terminal protein) plays in viral replication and pathogenesis are not well-characterized. Recent reports have identified that murine norovirus NS1, the N-terminal portion of NS1-2, antagonizes the interferon pathway, but studies of full-length NS1-2 or the NS2 domain remain limited (17–19). Recombinant expression of NS1-2 from feline calicivirus (FCV), murine norovirus (MNV), and HuNoV GII.4 shows that the protein traffics to the endoplasmic reticulum (ER), concentrates perinuclearly, colocalizes with the ER-resident protein calnexin, and contains C-terminal hydrophobic sequences (20–22). In contrast, Norwalk virus (GI.1) NS1-2 (p48) was primarily found in the Golgi apparatus and implicated in disrupting ER-to-Golgi trafficking (23, 24). The similarities in ER/Golgi membrane association and domain organization of NS1-2 from different viruses suggest that NS1-2 may have a conserved function among caliciviruses.

The ER, and to a lesser extent Golgi apparatus, are important intracellular calcium (Ca^2+^) storage organelles, with the ER Ca^2+^ concentration being up to 1 mM (25, 26). As a ubiquitous secondary messenger, Ca^2+^ is at the epicenter of many cellular processes and host machinery tightly regulates Ca^2+^ levels to ensure low (nM) cytoplasmic Ca^2+^ concentrations at cellular rest (26–32). Importantly, Ca^2+^ signaling regulates several aspects of viral life cycles including entry, genome replication, and release (30, 33–35). To exploit Ca^2+^ signaling, many viruses express an ion channel (i.e., viroporin) to dysregulate Ca^2+^ homeostasis in order to usurp Ca^2+^-dependent host processes (30, 36–39). The best characterized Ca^2+^ disrupting viroporins are the nonstructural protein 4 (NSP4) from rotavirus (RV) and the 2B nonstructural protein of enteroviruses (EV) and some other picornaviruses (36, 37, 40–44). Like all *bona fide* viroporins, NSP4 and 2B have canonical biophysical motifs, including being oligomeric, having an amphipathic α-helix that forms the pore, and a cluster of basic residues that facilitate membrane insertion (37–39, 42–44). While no study has specifically looked at whether caliciviruses dysregulate Ca^2+^ signaling or have a viroporin, they belong to the picornavirus-like superfamily of positive-sense RNA viruses, among which there is considerable positional homology of the cognate proteins of the nonstructural polyprotein (23, 45, 46). Within this rubric, the picornavirus 2AB region constitutes the positional homolog of the calicivirus NS1-2 protein, and several sequence motifs in the NS1 are conserved in the 2A protein of some picronaviruses (23). While no functional homology between EV 2B and the NS2 region of NS1-2 has been yet been identified, it is tempting to speculate that NS1-2 may have viroporin activity and dysregulate host Ca^2+^ signaling analogous to that of EV 2B proteins.

In this study, we investigated the role of Ca^2+^ signaling in TV replication and whether TV NS1-2 has viroporin activity that can dysregulate Ca^2+^ homeostasis. Using long-term live-cell Ca^2+^ imaging, we sought to determine whether TV infection causes aberrant Ca^2+^ signaling during infection and identify the cellular Ca^2+^ pools critical for the TV-induced Ca^2+^ signaling. Finally, we tested TV NS1-2 for viroporin activity and determined whether the putative NS1-2 viroporin domain caused aberrant Ca^2+^ signaling similar to TV infection.

## Methods

### Cell lines, GECI lentiviruses, and viruses

All experiments were performed in LLC-MK2 cells. Lentivirus packaging and recombinant protein expression for western blot lysate production was performed in HEK293FT cells (ATCC CRL-3216). Cell lines were grown in high-glucose DMEM (Sigma: D6429) containing 10% fetal bovine serum (FBS) (Corning lot no. 35010167) and antibiotic/antimycotic (Invitrogen), and maintained at 37°C with 5% CO2. Lentivirus packaging in HEK293FT cells was performed as previously described (41). Briefly, LLC-MK2 cells were transduced with a lentivirus vector encoding GCaMP6s 1-day post-seeding (∼85% confluency). We confirmed positive expression of GCaMP6s at 48-72 hours post-transduction, then passaged cells 1:2 and added hygromycin (100 μg/mL) for selection of the LLC-MK2-GCaMP6s cell lines, henceforth referred to as MK2-G6s. We determined GCaMP6s activity and dynamic range using thapsigargin (0.5μM). Tulane virus (TV) stocks were made in-house by infecting cells with MOI 0.01 and harvesting at ∼95% cytopathic effect (CPE). Virus titer was determined by plaque assay. Irradiated virus controls were made by gamma-irradiating TV stocks for 19 hours.

### Replication assays

LLC-MK2 cells were seeded at 125,000 cells/well in 24 well plates (Costar 3524, Corning) and inoculated next day with TV at MOI 1 for 1 hour. Inoculum was removed, and cell media was replaced containing different extracellular Ca^2+^ conditions (0 mM Ca^2+^, 4 mM Ca^2+^), intracellular Ca^2+^ chelator 50 μM BAPTA-AM, or the sarco/endoplasmic reticulum calcium ATPase (SERCA) blocker thapsigargin (TG). Ca^2+^-free DMEM was purchased from Gibco (Cat #21068-028). Standard high-glucose DMEM (Sigma) has 1.8 mM CaCl2, which we refer to as “2 mM Ca^2+^, and media with 4 mM Ca^2+^ was made by adding 2 mM CaCl2 to the standard high-glucose DMEM (Sigma). We maintained TV-infected cells under these conditions until the positive control (normal media) had ∼90% CPE. Progeny virus was harvested by 3 freeze/thaw cycles, the virus yield was determined by plaque assay. For plaque assays, cells are seeded at 75,000 cells/well in 24 well plates and 2-days post-seeding, the cells were inoculated for 1 hour with 10-fold serial dilutions of the sample. Then, we removed the inoculum and added the overlay. Overlays for plaque assays was made mixing equal parts 1.2% Avicel (FMC Corporation) and 2X DMEM (Gibco). Plaque assays were harvested at 72 hours; fixed and stained with crystal violet (3% solution) to visualize plaques. Titer is represented as plaque forming units per milliliter (PFU/mL).

### One-step growth curves

One step growth curves for TV were performed using a modified protocol from previous reports (11, 15). Briefly, LLC-MK2 cells were inoculated with TV at MOI 1 in serum-free DMEM (0% FBS DMEM). At 1-hour post-infection (HPI), the inoculum was removed and replaced with 0% FBS DMEM. Cells were harvested at 0, 4, 6, 8, 10, 12, 16, 20, 24, and 28 HPI and virus yield determined by plaque assay. Each biological replicate was performed in duplicate.

### Long-term Ca^2+^ imaging experiments

Calcium imaging experiments were set up by adapting a protocol detailed in previous reports (47). MK2-G6s cells were seeded at 78,500 cells/well in 15 μ-slide 8 well chambers (Ibidi, Germany) and infected the next day with TV at the indicated MOI. After one hour, the inoculum was removed and replaced with FluoroBrite DMEM (Gibco). For studies involving pharmacological compounds, the FluoroBrite was mixed with DMSO (0.1%, vehicle control) or the indicated pharmacological compounds dissolved in DMSO. Then the slide was mounted into a stage-top environmental chamber (Okolab H301-Mini) that maintained 37°C with humidity control and 5% CO2. Time lapse live-cell Ca^2+^ imaging experiments were conducted for from ∼2 HPI until ∼18-24 HPI on a Nikon TiE epifluorescence microscope using a SPECTRAX LED light source (Lumencor) and a 20X Plan Apo (NA 0.75) objective. Images were acquired at 1-2 images/minute. Images were acquired and analyzed using the NIS elements advanced research software package (Nikon). Prior to image analysis, background camera noise was subtracted from the images using an averaged file of 10 no-light camera images. Cells that underwent division during the imaging run were excluded from analysis. Intracellular Ca^2+^ signaling over time was quantified by calculating the number of Ca^2+^ spikes per cell. This was determined as follows: Raw fluorescence intensity values were measured in Nikon software, then exported to Microsoft Excel to normalize the fluorescence to the first image (F/F0). The Ca^2+^ spikes were calculated by subtracting each normalized fluorescence measurement from the previous measurement to determine the change in GCaMP6s fluorescence (ΔF) between each timepoint. Ca^2+^ signals with a ΔF magnitude of >5% were counted as Ca^2+^ spikes. For each condition testes, Ca^2+^ spikes in ≥30 cells were determined.

### Heatmap generation

To generate heatmaps of the normalized GCaMP6s fluorescence over time for long-term Ca^2+^ imaging experiments we used the TidyR (48) and ggplot2 (49) packages available through R studio. Normalized GCaMP6s data from Excel was used to create a R-compatible file (.csv) containing the normalized fluorescence and the acquisition time data for the dataset, and imported the file into R. We used the TidyR package to organize data into a format accessible by ggplot2. We then used ggplot2 to generate heatmaps.

### Prediction of viroporin motifs *in silico*

We used the Hydropathy Analysis program at the Transporter Classification Database to generate Kyte & Doolittle Hydropathy and Amphipathic moment plots to identify putative viroporin motifs within full-length TV NS1-2 (50). Secondary structure, membrane topology, and membrane integration predictions were performed using PSIPred prediction analysis suite (website: http://bioinf.cs.ucl.ac.uk/introduction/) (51). Helical wheel plots to identify clustered basic residues within the putative viroporin domain were generated using the PepWheel analysis program at Heliquest (website: http://heliquest.ipmc.cnrs.fr/) (52).

### Expression vectors

*E. coli* expression constructs for the lysis assay were generated via ligation-independent cloning (LIC) using the pET46-Ek/LIC kit (MilliporeSigma, Darmstadt, Germany). The pET46-Ek/LIC constructs all have an N-terminal 6x histidine tag. Mammalian expression vectors were generated by inserting c-myc tag and mRuby3 red fluorescent protein upstream of full length NS1-2 and then subcloning this into the pTagRFP-N vector in place of TagRFP (Epoch Life Sciences, Missouri City, TX). This construct will be referred to as RFP-NS1-2. The NS1-2(Δ176) and NS1-2(Δ157) truncation mutations in both bacterial and mammalian expression vectors were generated using the NEB Q5 site-directed mutagenesis kit (New England Biolabs, Ipswich, MA). All constructs were sequence-verified using universal primers specific to the construct backbone (GENEWIZ, South Plainfield, NJ). The mammalian expression vector for EV 2B was generated by cloning the 2B from enterovirus 71 upstream into pTagRFP-N and the construction of the NSP4-TagRFP expression vector was previously described (53).

### Transfection experiments

MK2-G6s cells were seeded in 15 μ-slide 8 well chambers (Ibidi, Germany) and at 85% confluency transfected with mammalian expression constructs in Opti-MEM (ThermoFisher) and Lipofectamine 2000 (Invitrogen). Transfection was optimized so cells received 400 ng plasmid DNA and 0.5 μL of Lipofectamine 2000 per well. 10 μM trichostatin A (TSA) was added from 1-3 hours post-transfection. TSA is a histone deacetylase (HDAC) inhibitor used to increase expression from the vectors (54–56). Time-lapse Ca^2+^ imaging was performed beginning 8 hours post-transfection to capture expression kinetics, and up to 24 hours post-transfection to measure changes in Ca^2+^ signaling during expression of the RFP-tagged proteins.

### Deconvolution microscopy

LLC-MK2 cells were seeded in 15 μ-slide 8 well chambers (Ibidi, Germany) and transfected one day prior to imaging. Cells were transfected with intracellular markers for the plasma membrane (LCK-GFP; Addgene plasmid #61099), endoplasmic reticulum (pLV-ER GFP; Addgene plasmid #80069), Golgi apparatus (pLV-Golgi GFP; Addgene plasmid #79809), and mitochondria (HyPer-dMito, Envrogen). Control wells received TagRFP (Envrogen), while experimental wells received either full length RFP-NS1-2, RFP-NS1-2(Δ157) or RFP-NS1-2(Δ176). Cells were imaged 24 hours post-transfection on the DeltaVision LIVE high resolution deconvolution microscope (GE Healthcare) using the 60X/1.4 Plan-Apo NA oil objective (Olympus), and acquired using a pco.edge sCMOS_5.5 camera. Images were acquired and deconvolved in SoftWoRx software. Upon deconvolving, images were further processed in FIJI (ImageJ) to adjust for brightness/contrast and pseudo-coloring (57).

### *E. coli* Lysis Assay

*E. coli* Lysis assays were performed as previously described (41). Briefly, pET46-Ek/LIC constructs of the TV NS1-2 full-length and truncation mutants were transformed into BL21(DE3)pLysS cells. Transformations were plated on LB + 1% glucose + 100μg/mL ampicillin, + 35μg/mL chloramphenicol and grown at 37°C overnight. Isolated colonies were picked next day and cultured overnight in liquid LB + 1% glucose, + 100μg/mL ampicillin, + 35μg/mL chloramphenicol at 37°C in an orbital shaker at 250 rpm. The next day, overnight cultures were subcultured by 1:100 dilution into 200 mL LB + 1% glucose, + 100μg/mL ampicillin, + 35μg/mL chloramphenicol. Subcultures were grown at 37°C in an orbital shaker at 250 rpm for ∼3 hours to an OD630 between 0.3-0.5, then induced with 1mM IPTG. Absorbance measurements at 630 nm (OD630) were taken every 10 minutes for 90 minutes and normalized to the induction OD630 to determine the percent growth or lysis over time after induction. Each experiment was performed ≥3 times. Protein expression was determined by SDS-PAGE using a 4-20% Tris-glycine gel (Bio-RAD, Hercules, CA) and western blot for the 6x histidine tag. An uninduced culture served as the negative control for viroporin activity and NS1-2 synthesis.

### Membrane association experiment

Membrane association experiments were performed using a modified protocol from previously reported experiments (36, 41). Lysed membranes from a 200 mL culture were centrifugated at 21,000 x g for 20 minutes and supernatants decanted. Pellets were resuspended in PBS and sonicated 3 times for 1 minute on ice. Total lysate was collected after sonication. Then membranes were pelleted by ultracentrifugation at 49,000 x g for 1 hour using a TLA-100.3 rotor in an Optima TL Ultracentrifuge (Beckman Coulter, Indianapolis, IN), and the supernatant was collected for the soluble fraction. Finally, the membrane fraction pellet was resuspended in PBS + 1% SDS to solubilize membrane proteins. Samples from the total lysate, soluble fraction, and membrane fraction analyzed by western blot.

### Production of TV and Vpg antisera

For the anti-TV antisera to detect VP1, adult male and female CD-1 mice (purchased from Center for Comparative Medicine, Baylor College of Medicine) were immunized 5 times with CsCl2-gradient-purified TV at 10 μg/dose in AddaVax adjuvant (InvivoGen). Immunizations were given at 3-week intervals. For the anti-Vpg antisera, adult Balb/C were immunized 3 times with 10-20 μg/dose purified Vpg expressed in *E. coli*. The priming dose was given in Freund’s complete adjuvant and the subsequent boosts were given in Freund’s incomplete adjuvant. All experiments were performed in accordance with the recommendations in the Guide for the Care and Use of Laboratory Animals of the National Institutes of Health.

### Immunoblot analysis

Samples were prepared using procedures adapted from (36). Briefly, samples were mixed with 5X sample buffer containing 2-mercaptonethanol and boiled for 10 minutes at 100^◦^C. Samples were then run on a 4-20% Tris-glycine gel (Bio-RAD, Hercules CA) and transferred onto a nitrocellulose membrane using the Transblot Turbo transfer system (BIO-RAD, Hercules, CA). To detect the bacterial constructs of NS1-2 and NSP4, we used the mouse α-HIS Tag monoclonal antibody at 1:1000 (Genscript, Piscataway, NJ). To detect mammalian expression constructs of NS1-2, we used the mouse α-c-Myc monoclonal (clone 9E10) antibody at 1:1000 (R&D Systems, MN). To detect TV structural protein VP1, we used the mouse α-TV polyclonal we made in-house by hyperimmunizing CD1 mice with purified TV particles. To detect TV nonstructural protein Vpg, we used the mouse α-Vpg polyclonal antibody made by hyperimmunizing mice with bacterially expressed and purified Vpg. For loading control of mammalian cell lysates, we used the mouse α-GAPDH at 1:3000 (Novus Biologicals, CO). We used alkaline phosphatase-conjugated goat α-mouse IgG at 1:2000 (Southern Biotech, Birmingham, AL), and visualized using alkaline phosphatase substrate [Tris-base, nitro blue tetrazolium (NBT), 5-bromo-4-chloro-3-indolyl phosphate (BCIP)].

### Statistical analysis

Statistical analyses were completed using GraphPad Prism (ver. 7.03). Data in this manuscript is presented as mean ± standard deviation. We performed columns statistics to collect descriptive statistics and to determine the normality of the datasets. We then used the unpaired Student’s t-test for datasets with a parametric distribution, or a Mann-Whitney test for datasets with a nonparametric distribution. Differences were determined statistically significant if the p < 0.05. Authors had access to the data for this manuscript, and all authors approved the final manuscript.

### Data availability

RConsole code for the heatmaps generated in this paper is available upon request.

## Results

### TV infection disrupts host calcium signaling kinetics in LLC-MK2 cells

Ca^2+^ is a ubiquitous secondary messenger and many enteric viruses (*e.g.*, RVs and EVs) require elevated cytosolic Ca^2+^ to facilitate replication (30, 36–39, 42, 43). To determine if TV causes aberrant Ca^2+^ signaling like other enteric viruses, we examined whether Ca^2+^ signaling dynamics changed during TV infection. We infected LLC-MK2 cells stably expressing GCaMP6s (MK2-G6s) with different infectious doses [multiplicity of infection (MOI) 1, 5, 10] or γ-irradiated inactivated TV and performed live-cell fluorescent microscopy during the infection. GCaMP6s is a GFP-based genetically encoded Ca^2+^ indicator that reports changes in cytosolic Ca^2+^ as an increase in fluorescence (58). TV-infected MK2-G6s cells show increased cytoplasmic Ca^2+^ levels (Fig. 1A) beginning at roughly 8 HPI (MOI 10) and progressing for the remainder of the infection (Supplemental Video S1). The observed increase in Ca^2+^ signaling coincides with the synthesis of TV nonstructural proteins, assessed by western blot using an α-Vpg antisera (Fig. 1B) and TV structural proteins, assessed by western blot using an α-TV antisera to detect VP1 (Fig. 1C), which show increased TV protein production between 8-12 HPI. Further, based on a one-step growth curve, the increased cytosolic Ca^2+^ also coincides with the onset of progeny virus production, which occurs between 6-8 HPI (Fig. 1D). The increases in cytosolic Ca^2+^ were dynamic during TV infection (Supplemental Video S2). In infected cells, we noted that changes in cytosolic Ca^2+^ occurred through an increased number of discrete Ca^2+^ signals, much like what we recently observed in RV-infected cells (Fig. 1E) (47). We refer to these high-amplitude, transient Ca^2+^ signals as “Ca^2+^ spikes” and quantitated the number of Ca^2+^ spikes per cell during infection. Compared to uninfected controls, TV-infected cells have significantly more Ca^2+^ spikes/cell, but cells inoculated with γ-irradiated TV did not exhibit increased Ca^2+^ signaling (Fig. 1F). Together, these data indicate that increased Ca^2+^ signaling requires replication competent virus and occurs at later during infection, well after entry has occurred. Additionally, Ca^2+^ signaling in infected cells increases in an infectious dose-dependent manner, saturating at MOI=5 (Fig. 1F). To visualize the aberrant Ca^2+^ signaling induced by TV, we generated heatmaps plotting normalized GCaMP6s fluorescence over time (Fig. 1G). Heatmap data shows an increased number and magnitude of Ca^2+^ signals and that cytosolic Ca^2+^ levels change earlier and more frequently throughout infection as infectious dose increases (Fig. 1G). The heatmaps also show that MK2-G6s cells inoculated with γ-irradiated TV do not have increased Ca^2+^ signaling compared to mock-inoculated cells (Fig. 1G), consistent with the lack of increased Ca^2+^ spikes (Fig. 1F). Taken together, these data suggest that, like other enteric viruses, TV disrupts host Ca^2+^ signaling kinetics during infection.

**Figure 1.**
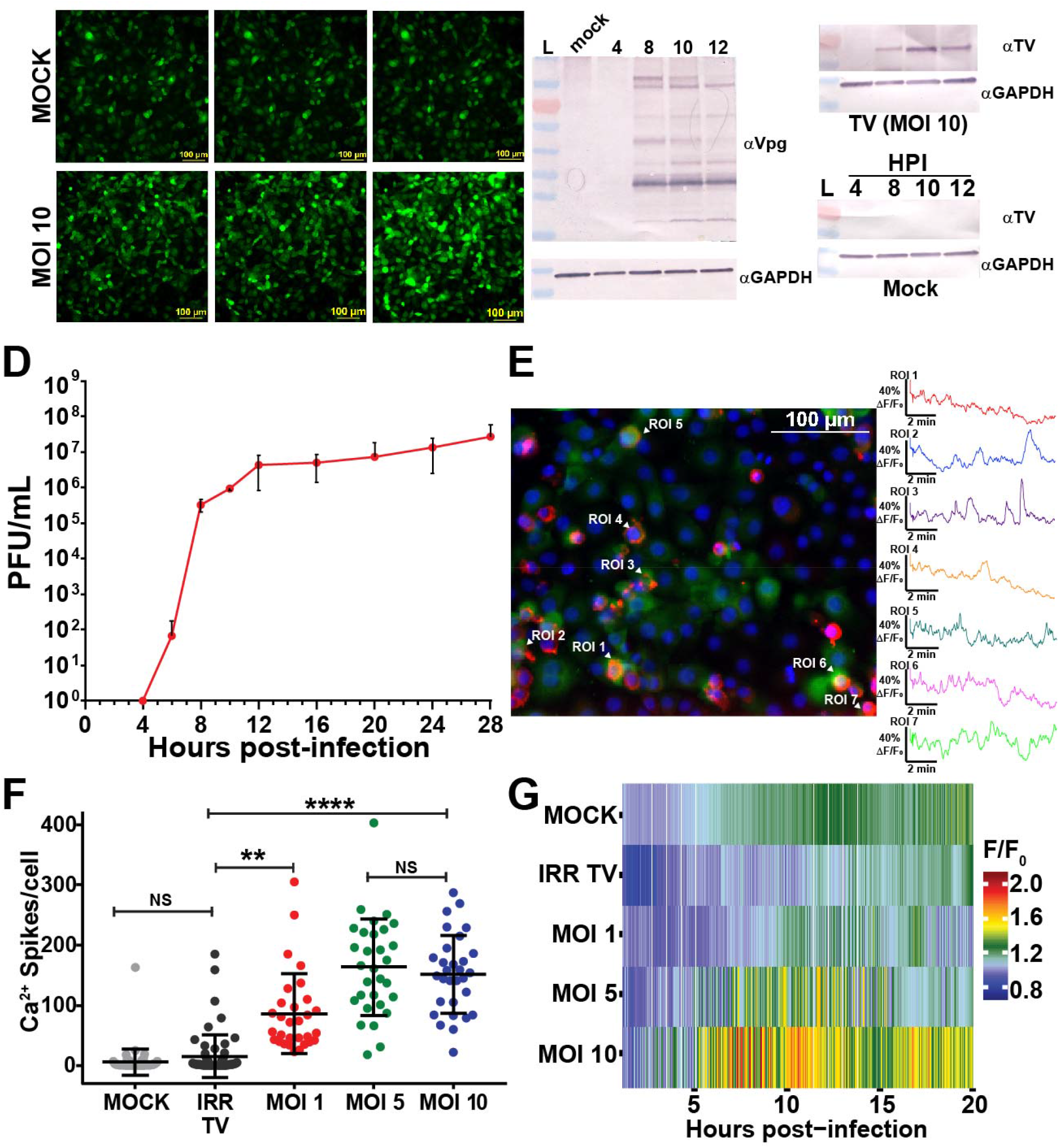
TV infection disrupts host calcium signaling kinetics in LLC-MK2 cells. **A**. Representative images at early (4 HPI), onset (8 HPI), and late (12 HPI) stages of mock (top) and TV-infected (bottom) LLC-MK2 GCaMP6s cells. **B, C.** Western blots for nonstructural protein Vpg (B) and structural protein VP1 (C) confirm that aberrant Ca^2+^ signaling in infected cells coincides with both structural and nonstructural protein synthesis. **D**. One-step growth curve for TV at a low MOI (1) shows that virus replication is concomitant with viral protein synthesis (1B, C) and with changes in Ca^2+^ signaling (1A). **E.** Still from overlay of α-Vpg staining (red) onto short (10 minute) continuous imaging runs of TV-infected cells (MOI=5) 12 HPI. Accompanying Ca^2+^ cell traces (right) shows the dynamic increases in cytosolic Ca^2+^ in infected cells **F.** Compared to mock, TV-infected cells have an increased number of Ca^2+^ spikes per cell that increases in an infectious dose-dependent manner, saturating at MOI=5. **G.** Heatmap data suggests that Ca^2+^ signaling increases with infectious dose, and that a higher MOI disrupts host Ca^2+^ signaling earlier in infection and sustains this aberrant Ca^2+^ signaling throughout. Mock and irradiated have similar heatmap profiles, suggesting that replication competent virus is required to drive these changes in Ca^2+^ signaling. Data shown as mean ± SD. **p<0.001, ****p<0.0001.

### Intracellular Ca^2+^ is critical for TV replication

Since we observed aberrant Ca^2+^ signaling during TV infection, we sought to determine whether Ca^2+^ was involved in TV replication. To test this, we manipulated extracellular and intracellular Ca^2+^ levels and determined their effect this had on TV yield. Doubling the extracellular Ca^2+^ concentration (∼4 mM) did not affect TV yield (Fig. 2A, right). In contrast, TV propagated in Ca^2+^-free media significantly reduced total yield (Fig. 2A, middle). Interestingly, plaques of TV propagated in Ca^2+^-free media were substantially smaller than that propagated in normal media, even though the plaque assay titrations were performed in normal media (Fig. 2C). Next, to investigate the role of intracellular Ca^2+^ during infection, we treated LLC-MK2 cells with BAPTA-AM, which chelates cytosolic Ca^2+^ and therefore buffers cytosolic Ca^2+^ (59, 60). TV replication in Ca^2+^-free media supplemented with BAPTA-AM (0Ca^2+^/BAPTA) was reduced up to 4-log (Fig. 2B), which was a greater inhibition than Ca^2+^-free media alone (Fig. 2A versus 2B). We next sought to determine whether intracellular Ca^2+^ stores are important for TV replication by testing the effect of thapsigargin (TG) on TV replication. TG is an inhibitor of sarco/endoplasmic reticulum Ca^2+^ ATPase (SERCA), which pumps cytosolic Ca^2+^ into the ER to help maintain ER Ca^2+^ stores. We treated TV-infected cells with TG and measured TV yield as above and found that TV replication is ∼3-log less in TG-treated than in DMSO-treated cells (Fig. 2B). Finally, we tested these different manipulations of extracellular or intracellular Ca^2+^ on TV yield a different time points during infection (8, 16, 24 HPI) (Fig. 2D). These studies confirmed that reduction of extracellular Ca^2+^ or treatment with TG significantly inhibited total virus replication; however, the rate of progeny virus production was not substantially reduced. Together, the replication assays demonstrate that intracellular Ca^2+^ levels facilitate TV replication, and that the ER Ca^2+^ store is particularly important for robust virus production.

**Figure 2:**
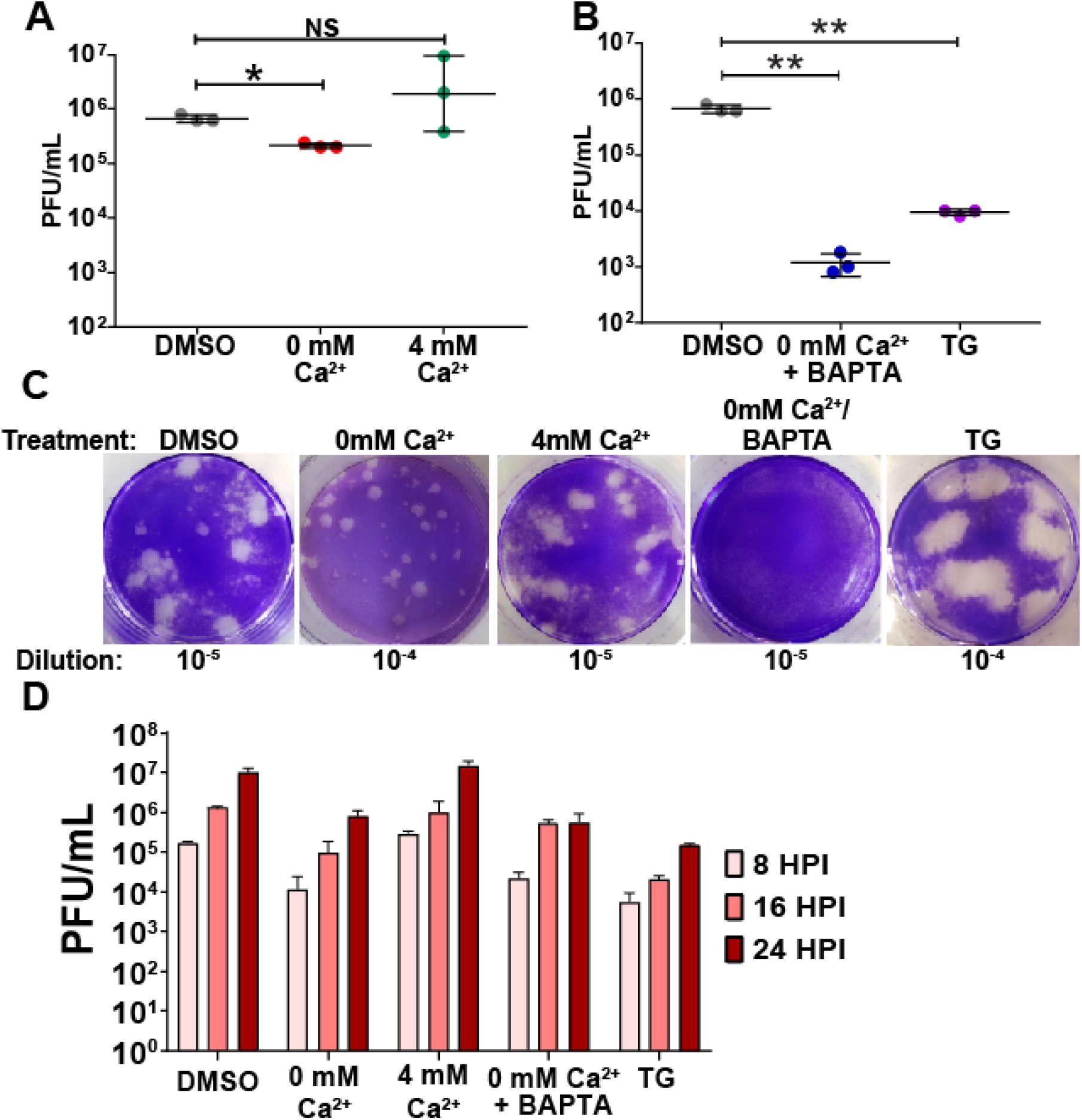
Intracellular calcium is critical for TV replication. **A.** Buffering out extracellular calcium hinders TV replication, significantly reducing the total plaque forming units (PFU). In contrast, excess extracellular Ca^2+^ (4 mM Ca^2+^, right) does not impact replication. **B.** Buffering intracellular calcium reduces replication. Depleting ER calcium stores with the SERCA-inhibitor thapsigargin and reducing cytoplasmic Ca^2+^ with BAPTA-AM significantly reduce TV infectious yield (PFU/mL). **C.** Representative images of plaques under normal (2mM) Ca^2+^ conditions (top, DMSO) and reduced Ca^2+^ (0 mM Ca^2+^/Bapta-AM, TG), conditions (bottom). **D.** Partial one-step growth curve data altering free intracellular (IC) and extracellular (EC) Ca^2+^. TV replication is stunted in Ca^2+^ free IC and EC conditions (0 mM Ca^2+^, 0 mM Ca^2+^/BAPTA-AM). Inhibiting ER Ca^2+^ replenishment with thapsigargin also blunts replication, suggesting that IC Ca^2+^ stores are critical for TV replication. Data shown as mean ± SD. *p<0.01, **p<0.001.

### TV-induced Ca^2+^ signaling requires ER Ca^2+^ stores

We next sought to determine the effect that the manipulations to extracellular and intracellular Ca^2+^ had on the TV-induced Ca^2+^ signaling exhibited in Figure 1. We altered extracellular and intracellular Ca^2+^ concentrations as above and performed live Ca^2+^ imaging of mock and TV-infected MK2-G6s cells. TV-infected cells in 2 mM Ca^2+^ (normal media) exhibited increased Ca^2+^ signaling, as observed above (Fig. 3A). Supplementing media with additional extracellular Ca^2+^ (4 mM Ca^2+^ total) did not further increase the Ca^2+^ spikes but removing extracellular Ca^2+^ abolished the TV-induced Ca^2+^ spikes (Fig. 3A). Using heatmaps, we plotted the relative change in GCaMP6s fluorescence over time and observed increased signaling starting at ∼8 HPI in both the 2 mM Ca^2+^ and 4 mM Ca^2+^ conditions (Fig. 3B). Further, the heatmaps show that infected cells in Ca^2+^-free media have a signaling profile that phenotypically mimics uninfected controls (Fig. 3B). Like the results obtained in replication assays, buffering cytoplasmic Ca^2+^ using BAPTA-AM reduced the number of Ca^2+^ spikes per cell to a level comparable to that of mock-infected cells (Fig. 3C) (Supplemental Video S2). Similarly, blocking the ER SERCA pump with TG significantly reduces TV-induced Ca^2+^ signaling (Fig. 3D), supporting replication data and demonstrating that ER Ca^2+^ stores are a critical source of Ca^2+^ for enhancing replication.

**Figure 3:**
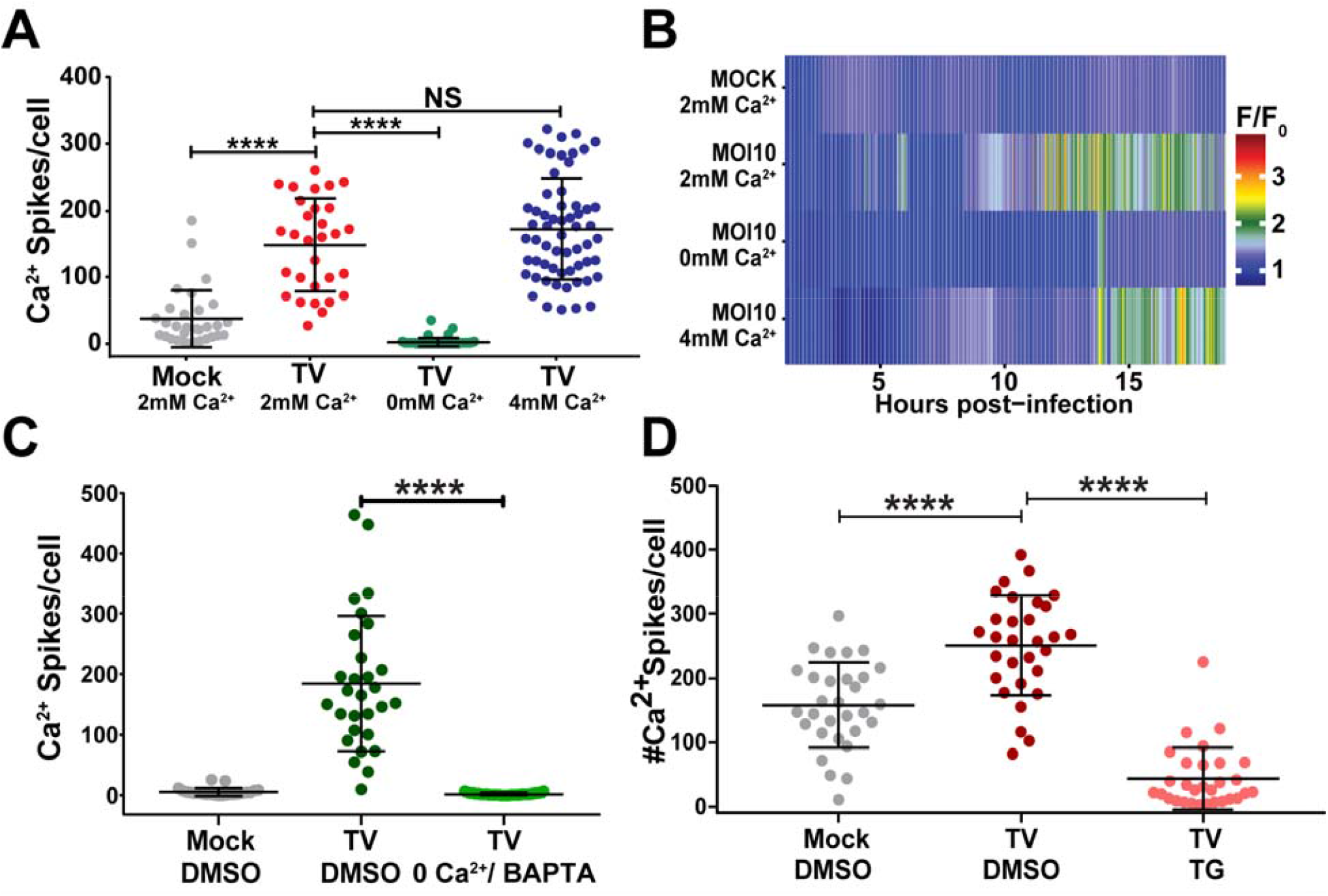
TV-induced Ca^2+^ signaling requires ER Ca^2+^ stores. **A**. Ca^2+^-free media reduces Ca^2+^ signaling in TV-infected cells, suggesting Ca^2+^ signaling is activated during infection. **B.** TV infection in 0 mM Ca^2+^ phenocopies mock Ca^2+^ traces in heatmap data, suggesting that EC Ca^2+^ facilitates TV infection. **C.** Intracellular Ca^2+^ chelator BAPTA-AM abrogated TV-induced Ca^2+^ signaling. BAPTA-AM-treated TV-infected cells (light green) returns Ca^2+^ signaling to uninfected levels (grey). **D.** Depleting ER Ca^2+^ with SERCA blocker thapsigargin (TG) significantly reduces TV-induced Ca^2+^ signaling (pink), suggesting that ER Ca^2+^ stores are a key source of Ca^2+^ leveraged during infection. Data shown as mean ± SD. ****p<0.0001.

### Tulane virus NS1-2 is targeted to the ER membrane

Our data indicates that TV activates aberrant Ca^2+^ signaling involving the ER Ca^2+^ store, much like the dysregulation of Ca^2+^ homeostasis by other enteric viruses observed in RV and EV infections. Both RV and EV encode a viroporin, or viral ion channel, that targets the ER Ca^2+^ store to activate aberrant Ca^2+^ signaling pathways that are critical for virus replication (35, 36, 38, 39, 42, 43). Viroporins are integral membrane proteins that have some common characteristics, including being oligomeric, having an amphipathic α-helix that serves as the channel lumen through the membrane, and a cluster of basic amino acid residues that facilitate insertion into the membrane (24, 34, 35, 39, 61). Previous work with NS1-2 from several different caliciviruses shows it is membrane-associated and primarily to the ER (17–20, 24) and/or Golgi (17, 20, 21, 23). Thus, we hypothesized that calicivirus NS1-2 could be a viroporin involved in the aberrant Ca^2+^ signaling we observed during TV infection. Notably, the calicivirus NS2 domain is the positional homolog of the EV 2B viroporin (Fig. S1). This is potentially significant because previous studies have found conserved functional characteristics between the positional homologs of the other nonstructural proteins (20, 21, 23, 46, 61–64), and functional homology between EV 2AB and human norovirus (HuNoV) GII.4 NS1-2 (20, 23). Additionally, when performing multiple sequence alignments of other calicivirus NS1-2s, we found that the C-terminal domain (CTD) of this protein is highly conserved (Fig. S2). To determine whether TV NS1-2 has viroporin-like characteristics, we examined TV NS1-2 for viroporin motifs. First, we performed a Kyte-Doolittle plot to detect hydrophobic regions of NS1-2 and an amphipathicity plot to identify potential amphipathic domains (Fig. 4A). We found that aa195-215 (Fig. 4A, dark green box) in the CTD of NS1-2 has a high amphipathic moment. We then used PSIPred (51) to model NS1-2 predicted secondary structure (Fig. 4B). Output from this analysis suggested that the NS1-2 CTD was predominantly comprised of alpha-helices (Fig. 4B, pink residues), and accompanying confidence scores for prediction of these C-terminal helices were ≥75% (Fig. S3). Interestingly, the region of peak amphipathicity (Fig. 4A) was located within one of the PSIPred helix predictions of the CTD (Fig. 4B, dark green bar) and contained clustered basic residues (blue asterisks), two key features of viroporins. Additionally, NS1-2 topology modeling identified two putative transmembrane domains (TMD): the first (TMD1) from aa164-179, and the second (TMD2) from aa202-225 (Fig. 4C, top). The membrane topology schematic indicated that both TMD1 and TMD2 had predicted pore-lining regions within their helices (Fig. 4B bottom left). To explore this, we used HeliQuest (52) to generate a helical wheel diagram for TMD2 (aa198-215), since TMD2 had the clustered basic residues common among viroporins. The helical wheel shows that TMD2 is highly amphipathic with clear polar and non-polar faces to the helix (Fig. 4D). The calculated hydrophobic moment for TMD2 is 0.522, supporting the above amphipathicity predictions (Fig. 4A). Given the results of these computational studies, we predicted that NS1-2 TMD2 (aa195-215) is a viroporin domain and set out to test this prediction experimentally.

**Figure 4:**
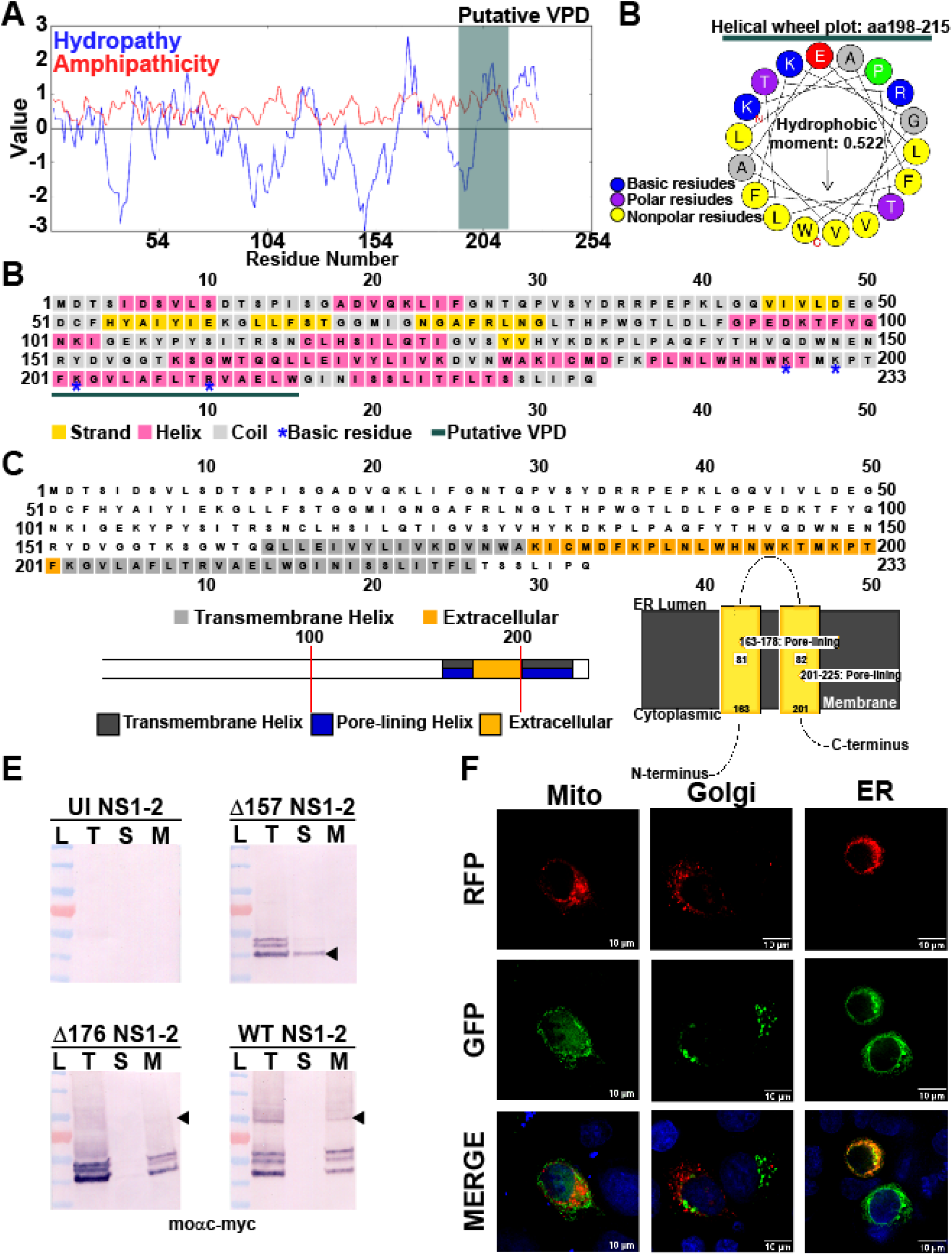
Tulane virus NS1-2 is targeted to the ER membrane. **A.** Predictive modeling of TV NS1-2 reveals that is has essential features of *bona fide* viroporins. Kyte-Doolittle-hydropathy plots predict an amphipathic moment from aa195-212 (dark green bar), consistent with alpha-helical structure required for channel formation. **B.** PSIPred secondary structure algorithms predict the C-terminus of NS1-2 is helical in nature, with the putative viroporin domain (VPD) contained to helices. **C.** PSIPred membrane topology predictions suggest that NS1-2 has two transmembrane helices (grey cubes, top figure). PSIPred algorithms predicting transmembrane helices suggest NS1-2 transmembrane domains are pore-lining (bottom left) and propose a model of membrane insertion and orientation where the putative VPD (aa195-212) comprises the pore-lining helix (bottom right). **D.** Helical wheel plot generated from the NS1-2 amphipathic segment (dark green bar) show clustered basic residues (green asterisks) and a hydrophobic moment of 0.522 from aa198-215, coinciding with the putative VPD **E.** Mammalian expressed full-length RFP-NS1-2 and RFP NS1-2 Δ176 is membrane-associated, but RFP NS1-2 Δ157 is not. Both the total fraction (T) and membrane pellets (M) extracted with 1% SDS contain RFP-NS1-2 and Δ176, but centrifuged supernatant (S) does not, suggesting that RFP-NS1-2 and Δ176 are membrane-associated proteins. In contrast, the supernatant contains RFP-NS1-2 Δ157. Further, immunoblot assays run under non-reducing conditions show that full-length RFP-NS1-2 and Δ176 oligomerize (black arrowheads) **F.** Co-transfection experiments using intracellular markers for predominant intracellular Ca^2+^ stores mitochondria (mito), Golgi, and endoplasmic reticulum (ER) to determine whether TV NS1-2 associated with any intracellular organelle(s). Based on deconvolution microscopy data, RFP-NS1-2 localized to the ER (right), but not with the Golgi (middle). RFP-NS1-2 did not localize to the mitochondria (left) (N≥2).

First, we tested whether TV NS1-2 was an integral membrane protein and whether it localized to the ER similar to NS1-2 from other caliciviruses. To do so, we generated bacterial and mammalian expression vectors of full-length NS1-2. For mammalian expression vectors, we N-terminally fused full-length NS1-2 to mRuby3 (henceforth referred to as RFP-NS1-2). From these constructs, we generated two truncation mutants of WT NS1-2 in both mammalian and bacterial expression vectors: the first, NS1-2 Δ176, was predicted to have TMD1 but lack the viroporin domain, and the second, NS1-2 Δ157, was predicted to lack both TMD1 and the VPD. We then transfected wild-type, full-length (WT) RFP-NS1-2, RFP-NS1-2 Δ157, and RFP-NS1-2 Δ176 into HEK 293FT cells and harvested cell suspensions next day. Samples following cell lysis, sonication, and fractionation were collected for SDS/PAGE western blots. We found both Δ176 and WT TV NS1-2 in the total fraction (T) and membrane pellets (M), but not in the supernatant (S), suggesting that TMD1 mediates membrane association (Fig. 4E). Additionally, in the non-reducing, unboiled conditions used, oligomers of both both Δ176 and WT RFP-NS1-2 were detected by western blot (Fig 4E, arrowheads). Similar results were obtained from membrane fractionation of analogous bacterially expressed NS1-2 constructs (Fig. S4). Using the mammalian expression vectors of RFP-NS1-2, we performed colocalization assays with fluorescent markers of the ER, Golgi apparatus, and mitochondria. RFP-NS1-2 showed no colocalization with the Golgi or mitochondria (Fig. 4F). In contrast, RFP-NS1-2 strongly colocalized with the ER-GFP marker (Fig. 4F), indicating that, like NS1-2 from other caliciviruses and EV 2B and RV NSP4, TV NS1-2 traffics to the ER membrane.

### TV NS1-2 has viroporin activity that disrupts Ca^2+^ signaling

Since our predictive modeling suggested that NS1-2 met the biophysical requirements for a viroporin, and our live-cell Ca^2+^ imaging data exhibited large changes in cytosolic Ca^2+^ during TV infection, we tested whether NS1-2 has viroporin activity. We performed the *E. coli* lysis assay, which is a classical viroporin functional assay, wherein viroporin expression by BL21(DE3)pLysS *E. coli* results in permeabilization of the inner membrane, resulting in T7 lysozyme-mediated cell lysis (41). This assay has been used to identify and initially characterize many viroporins (36, 65, 66). We expressed full-length HisNS1-2 in BL21(DE3)pLysS cells and measured optical density (OD) over time following protein induction with IPTG. For the lysis assay, strong viroporin activity is characterized by large decreases in OD over time, whereas no viroporin activity is characterized by increases in OD over time. Our results show that induced NS1-2 has strong viroporin activity, similar to that of RV NSP4, our positive control for viroporin activity (Fig. 5A). We see no changes in OD over time for uninduced NS1-2, indicating that HisNS1-2 viroporin activity correlated with protein expression, detected by immunoblot for the 6xHis-tag (Fig. 5B). We then asked whether recombinant expression of RFP-NS1-2 alone increases Ca^2+^ signaling in MK2-G6s cells. To test this, we transfected MK2-G6s cells with mammalian expression vectors for RFP-NS1-2 as well as RFP-NSP4 and RFP-EV 2B, our positive controls for viroporin-mediated Ca^2+^ signaling. Expressing RFP-tagged viroporins in MK2-G6s cells significantly increases both the number and amplitude of Ca^2+^ spikes. However, this was not observed in cells expressing RFP alone, as illustrated by the representative single-cell traces (Fig. 5C) (Supplemental Video S3). As above, we quantitated the number of Ca^2+^ spikes and confirmed that recombinant expression of RFP-NS1-2 increased the number of Ca^2+^ spikes per cell approximately 2-fold, similar to that of EV 2B and RV NSP4 (Fig. 5D). Taken together, our results demonstrate that TV NS1-2 has viroporin activity in the lysis assay, similar to *bona fide* viroporins, and causes aberrant host Ca^2+^ signaling when expressed in mammalian cells.

**Figure 5:**
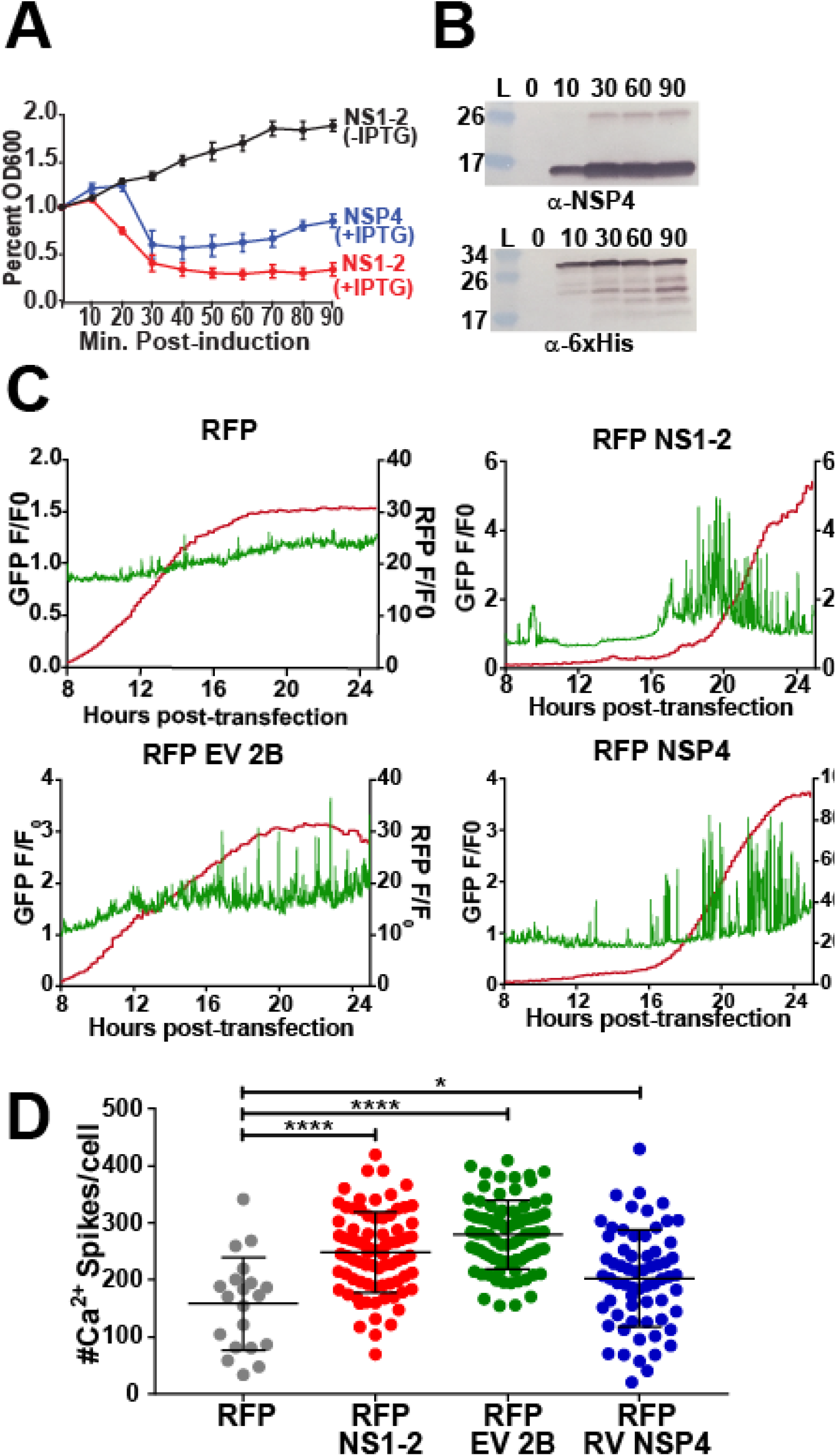
V NS1-2 has viroporin activity that disrupts Ca^2+^ signaling in mammalian cells. **A.** Inducing TV NS1-2 in the lysis assay strongly reduces optical density similar to rotavirus NSP4, the positive control for viroporin activity. B. Western blot data to verify protein expression during the lysis assay for TV NS1-2 (bottom) and RV NSP4 (top). **C, D.** Mammalian recombinant RFP-NS1-2 increases the number (B) and amplitude (C, top row, right) of Ca^2+^ spikes when transfected into cells similar to RV NSP4 and EV 2B, the viroporin controls for these experiments. Data shown as mean ± SD from ≥8 fields-of-view. *p<0.01, ****p<0.0001.

### NS1-2 viroporin activity maps to the putative viroporin domain

Our computational studies above identified a putative TV NS1-2 VPD from aa195-212. To determine whether the NS1-2 viroporin activity maps to this putative VPD, we generated C-terminal truncation mutants in bacterial expression vectors with deletions after aa212 (A212-Δ), after aa194 (W194-Δ), or after aa176 (D176-Δ) and characterized them in the lysis assay (Fig. 6A). We found that the A212-Δ truncation (red) had strong lysis activity comparable to full-length NS1-2 (black) (Fig. 6B). In contrast the D176-Δ truncation (blue) exhibited no lysis activity, comparable to uninduced NS1-2 (grey) (Fig. 6B). Immunoblot analysis confirm that protein expression correlated with viroporin activity, and that the impaired activity of W194-Δ was not due to lower expression levels, since the expression was comparable to that of full-length and A212-Δ (Fig 6C). Since the W194-Δ truncation (green) had impaired viroporin activity, this suggests that the VPD functionally extends to aa177-212.

**Figure 6:**
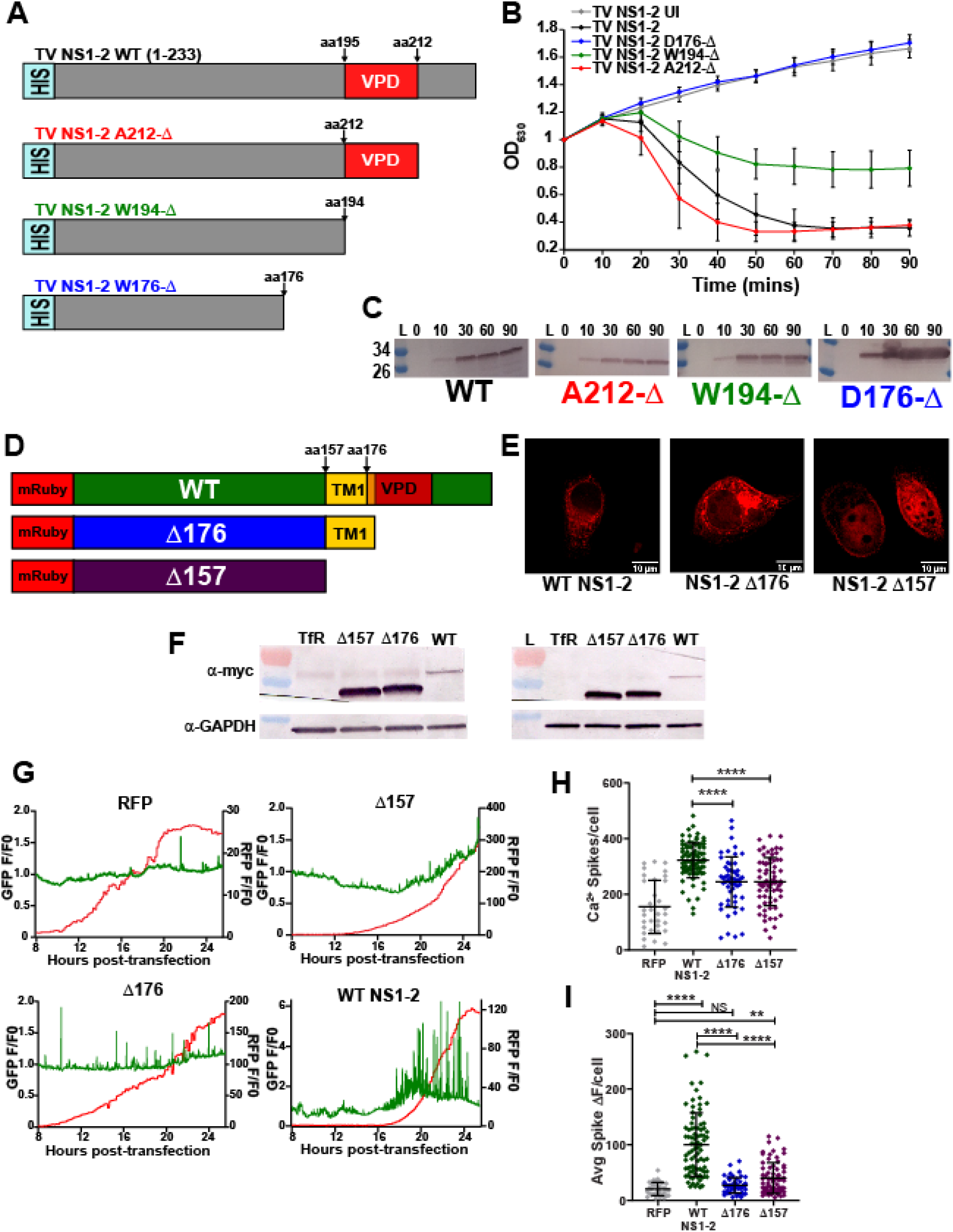
NS1-2 viroporins mutants do not increase cytoplasmic Ca^2+^. **A.** Schematic of TV NS1-2 C-terminal truncation mutants to functionally map the viroporin domain. **B.** In the lysis assay, truncating the C-terminal domain to amino acid 212 (red) results in wild-type activity (black), but truncating to W194 (green) impairs activity. Truncating to D176 (blue) abrogates viroporin activity, suggesting the viroporin domain functionally spans from aa177-212. **C**. Western blots verifying protein expression in the lysis assay. **D.** Schematic for the mammalian C-terminal truncation mutant constructs. **E.** IF data for truncation mutants. The Δ157 is cytoplasmic (far right), whereas the Δ176, which retains one transmembrane segment, is membrane localized (middle). **F.** Western blot data confirm the Δ157 and Δ176 mutant constructs. The left blot is run with 20 μL/well to visualize FL NS1-2, whereas the right blot is run with 5 μL/well to resolve the size difference between the Δ157 and 176 NS1-2 mutants. **G.** Both the Δ157 and Δ176 truncation mutants have significantly less Ca^2+^ spikes/cell compared to wild-type full length RFP-NS1-2. **H.** Compared to full length RFP-NS1-2, both the Δ157 and Δ176 truncation mutants have a significant reduced Ca^2+^ spike amplitudes, resulting in a change in cytosolic fluorescence (ΔF) that phenotypically mimics RFP alone. Data shown as mean ± SD from ≥8 fields-of-view. **p<0.001, ****p<0.0001.

Next, we characterized truncation mutants for their activation of aberrant Ca^2+^ signaling in MK2-G6s cells. Since recombinant expression of full-length RFP-NS1-2 induced aberrant Ca^2+^ signaling (Fig. 5D), we tested whether truncating the putative viroporin domain alone (Δ176) or both TMDs (Δ157) would compromise NS1-2-induced Ca^2+^ signaling (Fig. 6D). First, we examined the subcellular distribution and expression levels of the constructs. While the full-length and Δ176 truncation both appeared reticular, the Δ157 truncation had cytoplasmic distribution, consistent with it lacking both TMDs (Fig. 6E). Immunoblot analysis shows that the expression of both truncations was much greater than that of full-length NS1-2 (Fig. 6F, left blots) and by loading less lysate we can better resolve the 2 kDa size difference in the Δ157 and Δ176 truncations (Fig. 6F, right blots). Next, we examined whether these truncations could induce Ca^2+^ signaling by long-term live Ca^2+^ imaging in MK2-G6s cells. Individual cell traces illustrate that neither the Δ157 nor the Δ176 truncation dramatically increased Ca^2+^ signaling similar to full-length RFP-NS1-2 (Fig. 6G). Quantitation of the Ca^2+^ spikes per cell showed that while both truncations exhibited higher Ca^2+^ signaling than RFP alone (Fig 6G), the amplitude of these spikes was significantly reduced compared to full-length RFP-NS1-2 (Fig. 6H). The significant reduction in the number and amplitude of Ca^2+^ spikes/cell for both mutants highlights the critical importance of an intact VPD for disrupting host Ca^2+^ signaling. Together this work demonstrates that TV NS1-2 is an ER-targeted viroporin that induces aberrant Ca^2+^ signaling.

## Discussion

As obligate intracellular pathogens, viruses are adept at exploiting host pathways to facilitate replication. Viruses from many different taxonomic families activate aberrant Ca^2+^ signaling because Ca^2+^ signals are used by all cells to regulate a vast array of cellular functions. Therefore, this represents a powerful strategy to reconfigure host cell physiology via targeted disruption of host Ca^2+^ homeostasis. The overarching goal of this study was to determine whether dysregulation of Ca^2+^ signaling is a characteristic of caliciviruses and if this is due to the production of a viroporin protein similar to picornaviruses. To address these questions, we studied TV, as a model calicivirus, using a combination of live-cell Ca^2+^ imaging and other classical techniques. The major new findings of this study are as follows: (i) TV infection causes aberrant Ca^2+^ signaling that coincides with viral protein synthesis and replication, (ii) cellular Ca^2+^ is critical for TV replication and buffering of cytosolic Ca^2+^ severely reduced viral yield, (iii) TV NS1-2 has viroporin activity and dysregulates Ca^2+^ signaling in mammalian cells similar to TV infection, and (iv) NS1-2 viroporin activity maps to a C-terminal integral membrane viroporin domain and truncation of this domain abrogates the NS1-2-induced activation of Ca^2+^ signaling. To our knowledge, these results are the first to show exploitation of Ca^2+^ signaling by a calicivirus and identification of NS1-2 as a Ca^2+^-disrupting viroporin. These findings further extend the functional homology between the calicivirus nonstructural proteins and their picornavirus positional homologs.

The exploitation of host Ca^2+^ signaling to facilitate virus replication is a common feature of many viruses (30). Our finding that TV coopts Ca^2+^ signaling is consistent with previous studies showing that elevated Ca^2+^ levels are important for picornavirus replication, especially since caliciviruses and picornaviruses utilize a similar replication strategy (43). Similar to other Ca^2+^-disrupting viruses, TV also induces aberrant Ca^2+^ signaling peak of virus replication, many hours after cell entry. This is consistent with the reduced virus yield in media with reduced extracellular Ca^2+^ or treatments to buffer cytosolic Ca^2+^ (BAPTA-AM) or block refilling of ER Ca^2+^ stores (TG). Further, as we recently reported for RV infection, the TV-induced increase in cytosolic Ca^2+^ manifests as many discrete Ca^2+^ signals rather than a monophasic increase in Ca^2+^ over the infection (47). This raises the question of what cellular pathways are activated by this Ca^2+^ signaling and how they benefit TV replication. Both RV and EV have been shown to exploit Ca^2+^ signaling to activate the biosynthetic early stages of autophagy, which facilitates virus replication through rearrangement of cellular membranes to form replication complexes (67). MNV infection of primary macrophages or RAW264.7 cell line activates autophagy, but, in contrast to RV and EV, autophagy limits MNV replication (68). Thus, it remains to be determined whether autophagy plays a role in calicivirus replication complex assembly or whether Ca^2+^ signaling regulates autophagy activation during calicivirus infection. Further, elevated Ca^2+^ signaling may serve to modulate cellular apoptotic responses. Strong monophasic increases in cytosolic Ca^2+^ activate apoptosis through mitochondrial Ca^2+^ overload, but transient and oscillatory Ca^2+^ fluxes serve as pro-survival signals (69). Activation of apoptosis has been seen in norovirus and feline calicivirus-infected cells, and caspase activation is critical for cleavage and release of MNV NS1 from NS1-2, which in turn modulates cellular innate immune responses (19, 22). Thus, increased transient Ca^2+^ signaling may serve to counteract apoptosis activation and help prolong cell viability to maximize virus replication.

Within the superfamily of picornavirus-like positive sense RNA viruses, there is positional homology between the ORF1 nonstructural proteins of caliciviruses (and likely astroviruses) and the P2-P3 nonstructural proteins of picronaviruses (23, 45, 46). We used this framework to determine whether TV NS1-2 exhibited viroporin activity, since the positional homolog, the picornavirus 2B protein, is a well-established Ca^2+^-conducting viroporin (34, 44). We found that TV NS1-2 has viroporin activity, similar to 2B and RV NSP4, and the viroporin activity mapped to the integral membrane NS2 domains. Since both the N-and C-termini are likely oriented in the cytoplasm, NS1-2 is classified as a Class II2 viroporin, similar to the picornavirus 2B proteins (34, 44). This topology is supported by the cytosolic accessibility of the NS1 domain and the need for the C-terminus to also be localized in the cytosol to enable cleavage by the NS6 protease. This raises the question of whether NS1-2 viroporin activity is conserved throughout the *Caliciviridae* family. Though 2B and NS1-2 lack appreciable primary sequence homology, this is not surprising because viroporins, even from the same virus family, often only share the common viroporin motifs (*i.e.*, having [i] an amphipathic α-helix, [ii] a cluster of basic residues, and [iii] being oligomeric) (33, 38, 39). Among NS1-2 from different caliciviruses, we found that these characteristic features are conserved, so we predict viroporin activity of NS1-2 is a common function. Furthermore, since blunting cytosolic Ca^2+^ signaling with BAPTA-AM reduced TV replication, blocking NS1-2 viroporin activity with mutations or drugs should also reduce replication. This is supported by a previous study showing that recombinant coxsackie B3 virus with mutations of the 2B viroporin exhibited significantly impaired replication or were completely replication deficient (70). Analogous studies can be done using the TV reverse genetics system once residues critical for viroporin activity can be identified through mutagenesis screens of the TV NS1-2 viroporin domain we mapped in this study.

The increased Ca^2+^ signaling observed in TV-infected cells is phenotypically similar to that induced by recombinant expression of full-length NS1-2, but the Ca^2+^ signaling is abrogated by truncation of the viroporin domain. Further, NS1-2 primarily localized to the ER, which is a major intracellular Ca^2+^ storage organelle. Thus, our model predicts that NS1-2 directly releases Ca^2+^ from the ER; however, it is likely that both NS1-2 and activation of host Ca^2+^ signaling pathways contribute to the observed Ca^2+^ signals. Ca^2+^ signals from NS1-2 require it to directly conduct Ca^2+^ and have a high enough conductance that the ER Ca^2+^ release event can be detected by a fluorescent Ca^2+^ indicator, yet these unitary events are challenging to detect even for large channels like the IP3R (71). Future studies using patch clamp electrophysiology are needed to confirm that NS1-2 conducts Ca^2+^ and determine its conductivity. Nevertheless, based on the similarities between NS1-2 and other Ca^2+^-conducting viroporins, EV 2B and RV NSP4, NS1-2 viroporin activity would reduce ER Ca^2+^ levels and this in turn will activate host Ca^2+^ signaling pathways. First, the moderately increased steady-state cytosolic Ca^2+^ levels could foster more ER Ca^2+^ release by potentiating the IP3R Ca^2+^ release channel (72). Second, reduced ER Ca^2+^ levels activate the store-operated Ca^2+^ entry (SOCE) pathway, wherein decreased ER Ca^2+^ levels activate the ER Ca^2+^ sensing protein stromal interaction molecule 1 (STIM1). Activated STIM1 translocates to ER microdomains adjacent to the plasma membrane and opens Ca^2+^ influx channels, like Orai1, to elevate cytosolic Ca^2+^ (31, 32). This Ca^2+^ influx, in concert with SERCA, helps to refill ER stores for continued signaling.

HuNoV and human sapoviruses cause outbreaks of AGE and are a major cause of foodborne illnesses. However, the molecular mechanisms of how these caliciviruses cause vomiting and diarrhea, the chief symptoms of AGE, have not been characterized. The dysregulation of Ca^2+^ signaling by TV may provide insights into the pathophysiology of enteric caliciviruses. Both IP3-mediated ER Ca^2+^ release and SOCE have been shown to activate chloride secretion from epithelial cells (73, 74). Additionally, in studies of other viroporins, the viroporin-induced elevated cytosolic Ca^2+^ induces cytoskeleton rearrangement, leading to disassembly of tight junctions and loss of barrier integrity (39). Hyperactivation of chloride secretion and loss of tight junctions would contribute to excess fluid secretion and diarrhea. In our study we have shown that dysregulated Ca^2+^ signaling is a feature of calicivirus infection using TV. Thus, future studies can further examine the role of aberrant Ca^2+^ signaling in calicivirus pathophysiology using human intestinal enteroid cultures that support the replication of many HuNoV strains (4).

In summary, we have shown that TV activates aberrant Ca^2+^ singling during infection, and cellular Ca^2+^ is critical for robust TV replication. Further, we found that the NS2 domain of the NS1-2 nonstructural protein is a viroporin that alone induces Ca^2+^ signaling similar to TV infection. Together, these results indicate that NS1-2 is functionally analogous to EV 2B and RV NSP4. While little is known about the function(s) of NS1-2, and particularly the NS2 domain of NS1-2, the similarity with other Ca^2+^ conducting viroporins may provide a broader insight for understanding NS1-2 functions. Finally, antiviral drugs against viroporins have been developed for influenza M2 and HIV Vpu (34). Thus, the NS1-2 viroporin may be a viable antiviral drug target against caliciviruses.

## Acknowledgments

This work was supported in part by NIH grants R01DK115507 (PI: J. M. Hyser) and R21AI137710 (CoPI: J. M. Hyser and T. Farkas). Trainee support for A.C.G. was provided by NIH grants F30DK112563 (PI: A.Chang-Graham) and the BCM Medical Scientist Training Program and support for both A.C.G. and A.C.S was provided by the Integrative Molecular and Biomedical Sciences Graduate Program (T32GM008231, PI: D. Nelson). Funding support for the BCM Integrated Microscopy Core includes the NIH (DK56338, CA125123), CPRIT (RP150578, RP170719), the Dan L. Duncan Comprehensive Cancer Center, and the John S. Dunn Gulf Coast Consortium for Chemical Genomics. We would like to thank Drs. Michael Mancini and Fabio Stossi for deconvolution microscopy assistance.

---

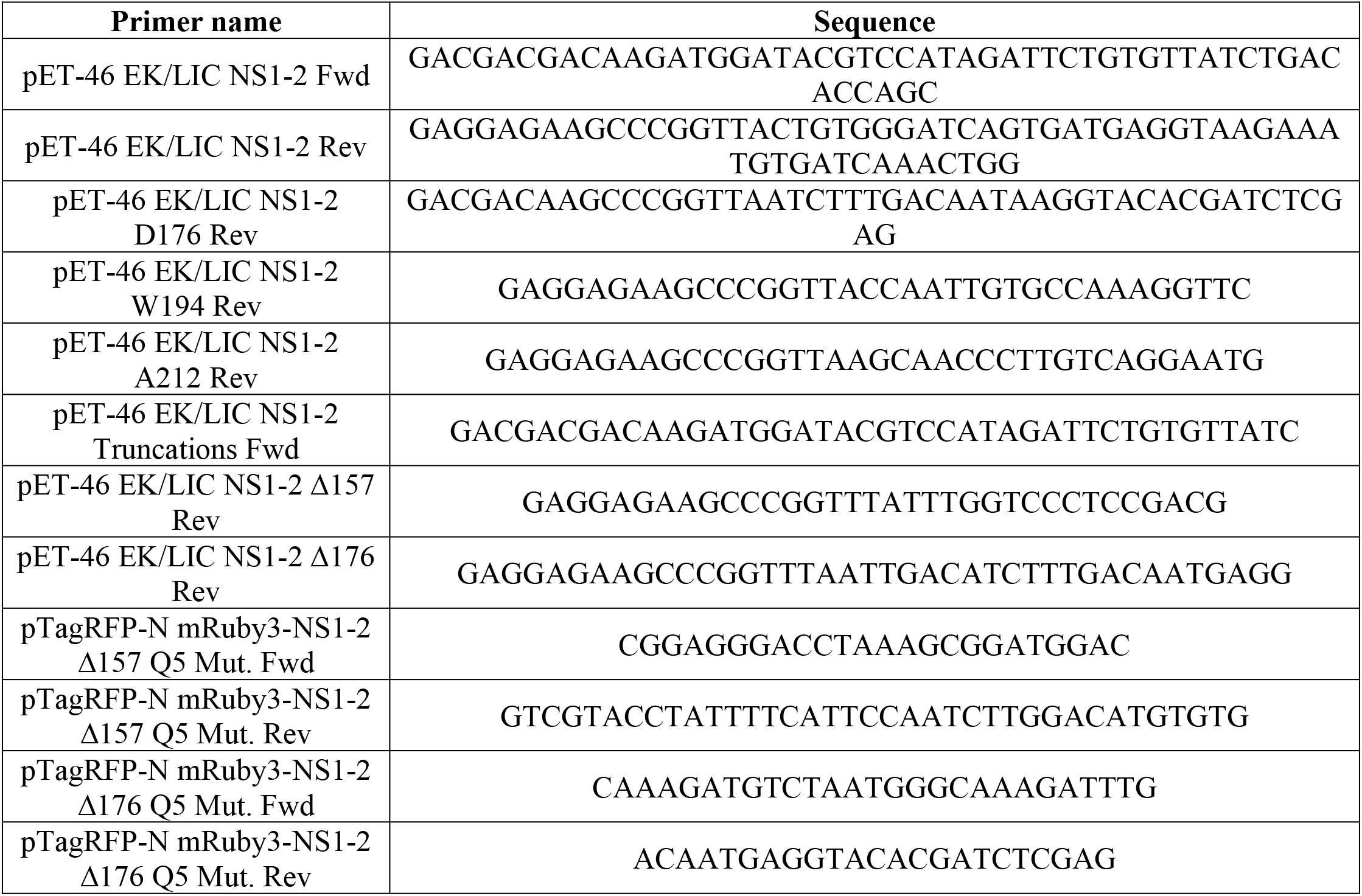
Primers:

## Notes

No conflicts of interest exist.

