## Supplemental Figures and Video legends for "Recovirus NS1-2 has viroporin activity that induces aberrant cellular calcium signaling to facilitate virus replication"

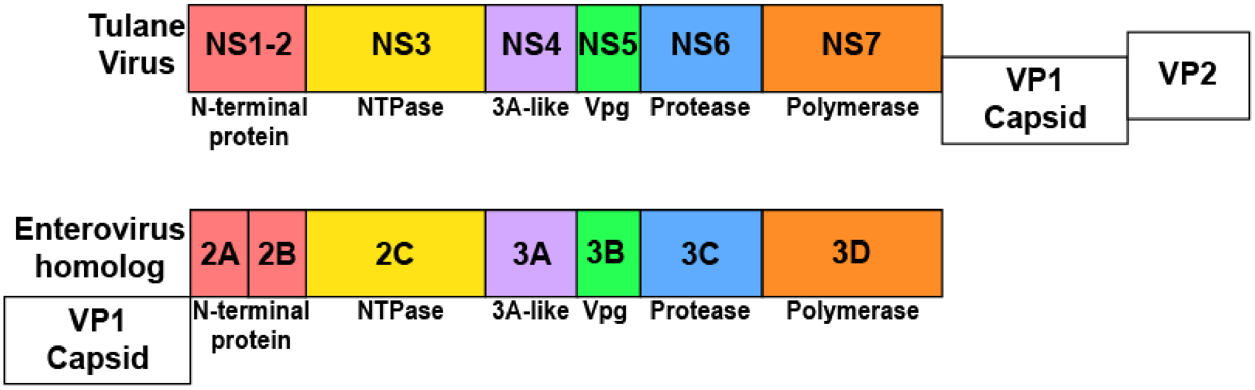


**Supplementary figure 1:** Aligning the nonstructural regions of both the TV and EV genome reveals that TV NS1-2 is the positional homolog of enterovirus (EV) 2B, a *bona fide* viroporin.

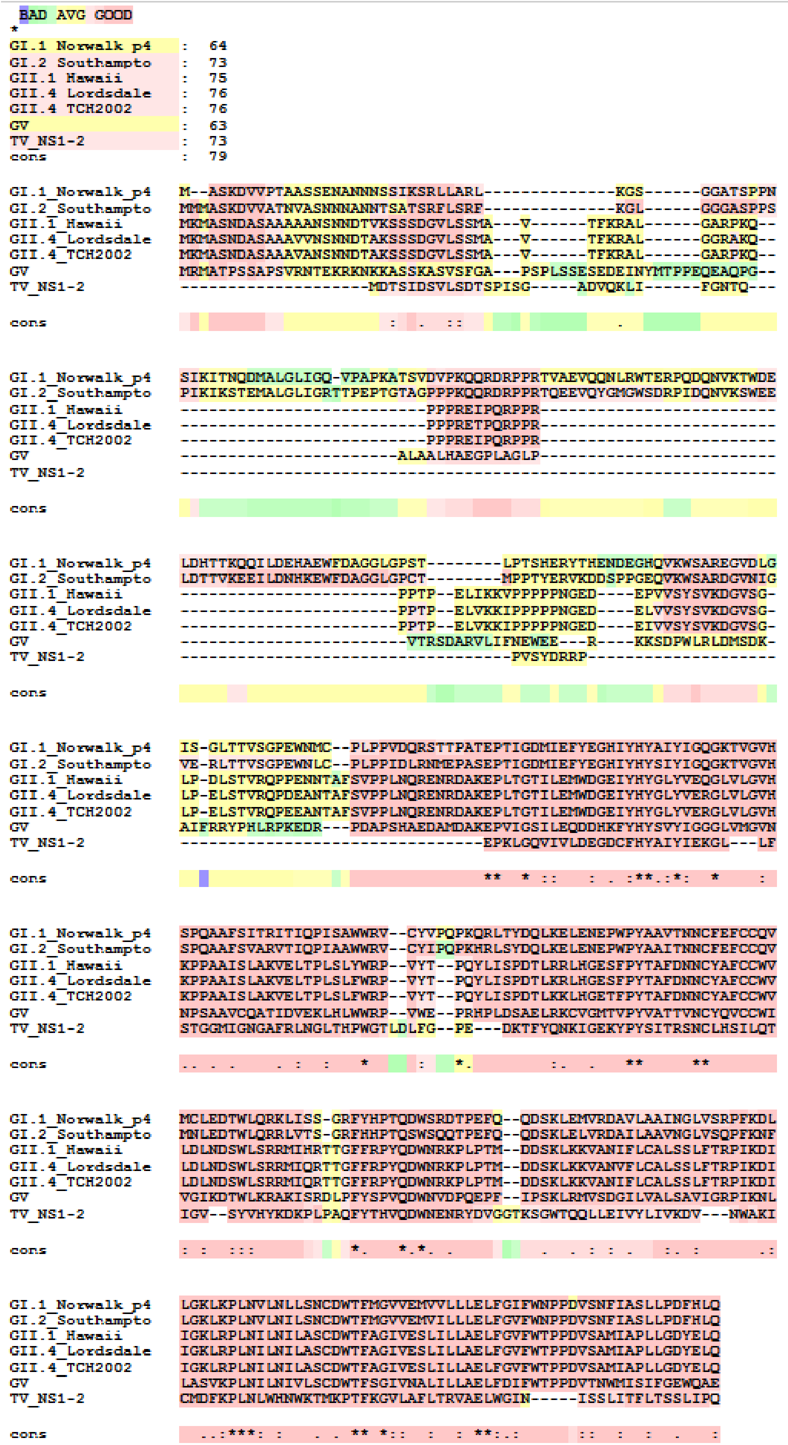

**Supplemental figure 2:** Multiple sequence alignment results of NS1-2 sequences from various norovirus strains and Tulane virus. Sequence alignment results through T-Coffee (71) show that the C-terminal domain (CTD) of NS1-2 across multiple noroviruses and tulane virus is well-conserved, including key basic residues at the same position within the CTD, which is necessary for biophysical conservation of viroporin activity.

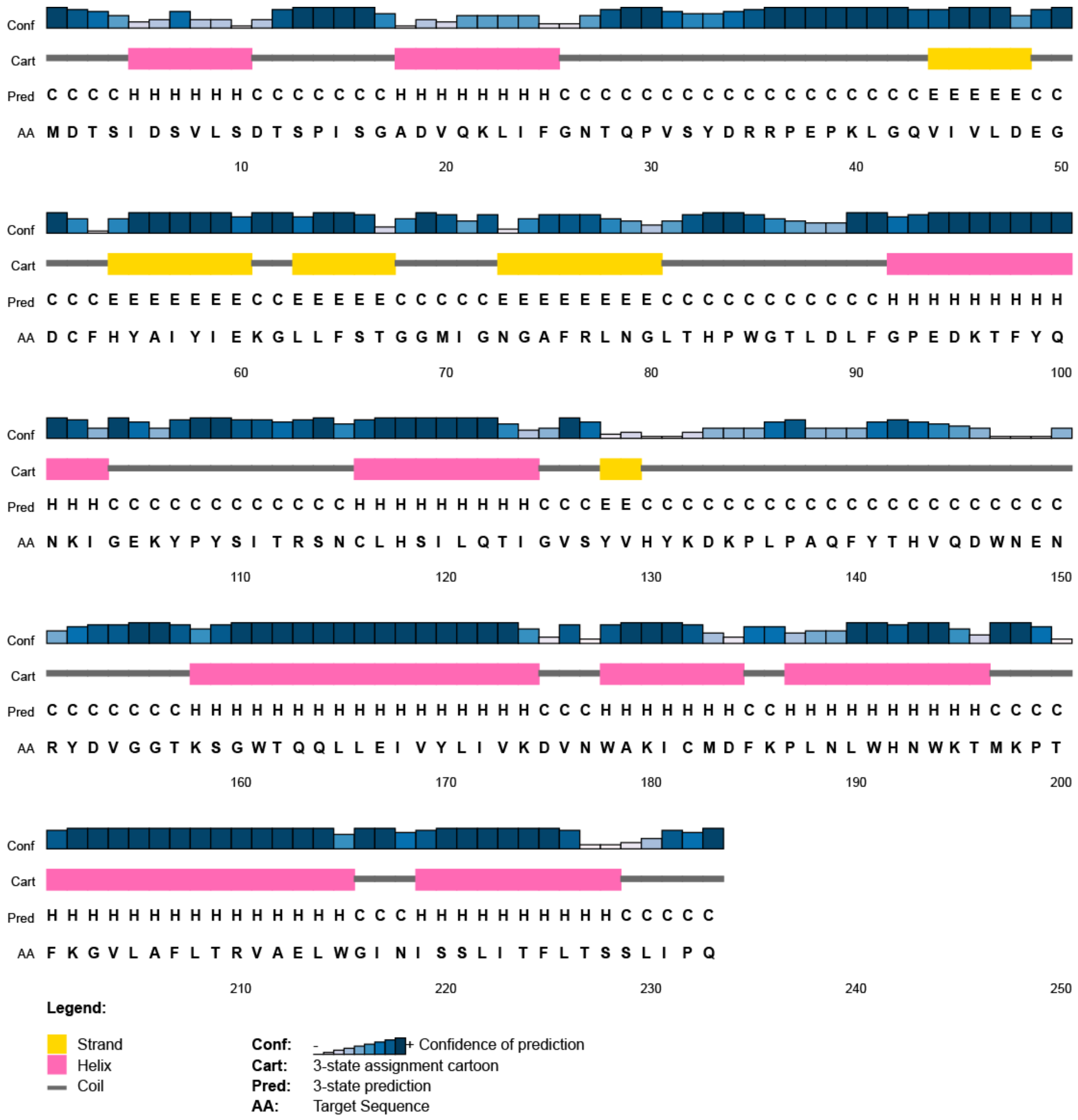

**Supplementary figure 3:** Confidence of prediction values of secondary structure for NS1-2 amino acid residues. Computational prediction for the viroporin domain (VPD) of NS1-2 places it between aa195- 215, which is predominantly helical, with ≥75% confidence of prediction throughout most of the C-terminal domain (aa 160-232, dark blue bars = conf).

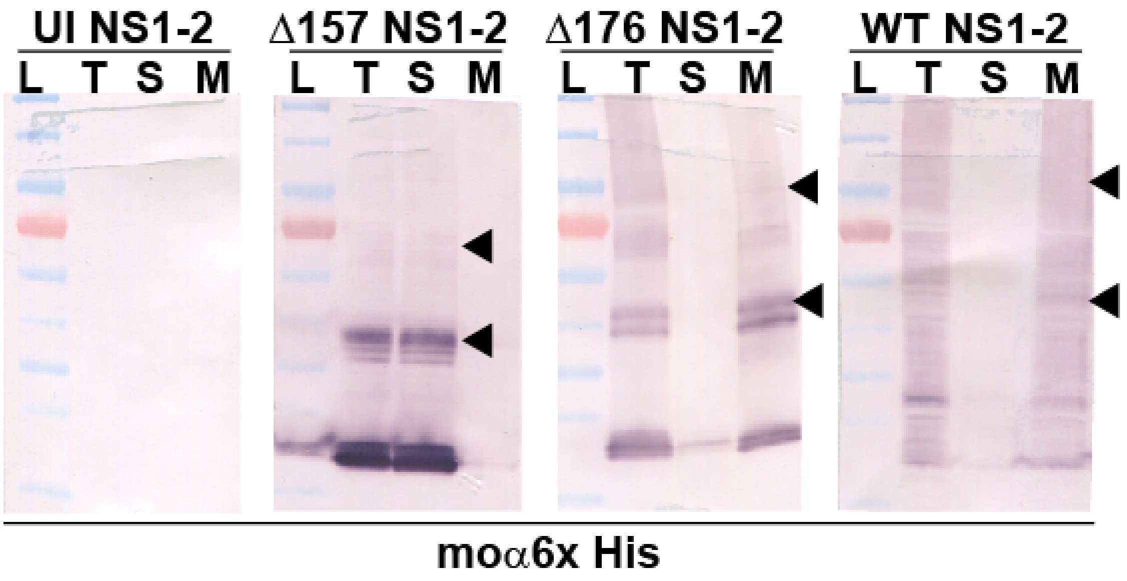


**Supplementary figure 4:** Immunoblot analysis of cell fractionation studies of bacterially expressed full-length WT NS1-2 and the Δ176 and Δ157 truncations. Samples of total cell lysate (T), the soluble fraction (S), or the membrane fraction (M) were resolved by SDS-PAGE and detected by immunoblot using an α-6xHis antibody. Oligomers of NS1-2 and the truncations are indicated by arrowheads.

**Supplementary Video 1:** Representative imaging run of LLC-MK2 GCaMP6s cells infected at a wide infectious dose (MOI 1, 5, 10) compared to irradiated TV. Movies qualitatively show that irradiated virus does not induce aberrant $Ca^{2+}$ signaling, but replication competent virus does. Quantification of this imaging run can be seen in figure 2.

**Supplementary Video 2:** IF staining for TV nonstructural protein α-Vpg overlaid onto short, continuous imaging run of TV-infected cells (MOI=5) 12 HPI. TV-infected cells (red) exhibit dynamic changes in cytosolic $Ca^{2+}$. Changes manifest as discrete $Ca^{2+}$ spikes which can be observed as large changes in cytosolic fluorescence.

**Supplementary Video 3:** Representative imaging run of LLC-MK2 GCaMP6s cells infected with TV (MOI 10, middle) compared to TV-infected cells treated with $0Ca^{2+}$/BAPTA (right). BAPTA treatment during TV infection abrogates TV-induced $Ca^{2+}$ signaling and results in a $Ca^{2+}$ signaling profile that mirrors mock LLC-MK2 cells (left).

**Supplementary Video 4:** Representative signaling from viroporin transfection experiments in LLC-MK2 GCaMP6s cells. Movies show that cells exhibit aberrant $Ca^{2+}$ signaling upon expression of RFP-tagged viroporins. TV NS1-2 (left) induces aberrant $Ca^{2+}$ signaling in cells with similar activity to RV NSP4 (middle) and EV 2B (right).
